# PEMapper / PECaller: A simplified approach to whole-genome sequencing

**DOI:** 10.1101/076968

**Authors:** H Richard Johnston, Pankaj Chopra, Thomas S Wingo, Viren Patel, Internation Consortium on Brain and Behavior in 22q11.2 Deletion Syndrome, Michael P Epstein, Jennifer Mulle, Stephen T Warren, Michael E Zwick, David J. Cutler

## Abstract

The analysis of human whole-genome sequencing data presents significant computational challenges. The sheer size of datasets places an enormous burden on computational, disk array, and network resources. Here we present an integrated computational package, PEMapper/PECaller, that was designed specifically to minimize the burden on networks and disk arrays, create output files that are minimal in size, and run in a highly computationally efficient way, with the single goal of enabling whole-genome sequencing at scale. In addition to improved computational efficiency, we implement a novel statistical framework that allows for a base-by-base error model, allowing this package to perform as well or better than the widely used Genome Analysis Toolkit (GATK) in all key measures of performance on human whole-genome sequences.

## INTRODUCTION

Whole-genome sequencing (WGS) using short reads on the Illumina platform is an increasingly cost effective approach for identifying genetic variation, with growing potential for both research and clinical applications^1,2,3,4^. A critical challenge lies in the development of efficient algorithms capable of rapidly and accurately identifying variable sites from among the enormous collection of sequence reads^5^. Given the large size of eukaryotic genomes, even modest false-positive or false-negative error rates can act as barriers to the success of genetic studies, and would inhibit the utility of such studies for both research and clinical applications.

The *de facto* standard methodology for mapping and calling variants is the so-called BWA/GATK Best Practices pipeline^6^, which was devised and validated for whole exome experiments and has greatly facilitated whole-exome studies for identification of disease causing variants^7-9^. While this pipeline can be used successfully at whole-genome scales, there are barriers to its use, particularly as the number of samples increases. BWA^10^, Bowtie^11^, and most other commonly used read mapping software packages are designed to run in low-memory footprints (i.e. less than 8 or 16 GB of RAM). Since whole-genome datasets are large (necessarily greater than 100 GB uncompressed for 30x coverage), these read mappers must continuously read and write large quantities of data to and from the disk. Sorting reads, in particular, is highly disk input/output (I/O) intensive. While a high-performance disk array can provide the needed I/O performance for a single instance of BWA/GATK processing^6^, no disk array can possibly accommodate the I/O performance required to run multiple GATK instances simultaneously on parallel processors. Moreover, even if the disk array itself could meet the demand, the network/fiber interconnects between the array and the computational nodes quickly become saturated. Simply put, while BWA/GATK Best Practices does an excellent job in a non-clustered environment, the “network cost” in a clustered environment significantly limits its performance for large whole-genome sequencing datasets.

GATK Best Practices has additional limitations. First, output files can be quite large. BAM files, required to store sequence alignment data, are almost always larger than the initial fastq files of nucleotide sequences, and Haplotype Caller output can be nearly half the size of the BAM files. Thus, total storage requirements to run the pipeline can approach 300GB, compressed, per sample for WGS data. Second, variant calling begins with individual samples (not collections of samples, *i.e.* joint calling), and as a result the distinction between sites called as homozygous reference genotypes and those called as missing (insufficient evidence to make a call), is not always maintained. Third, the GATK Best Practices joint genotyping caller, required to generate the highest quality genotype calls, does not scale well to whole-genome data. As currently implemented, the joint caller simply will not run on whole genome size files in sample collections larger than 10-20 human genomes, even on computers with 512GB of RAM. This seriously limits the utility of GATK for large-scale sequencing. Finally, the entire GATK Best Practices pipeline relies upon and uses enormous quantities of “previous knowledge” about the position and frequency of SNPs (Single Nucleotide Polymorphism) and indels (Insertion/Deletion Variants). This is both a strength, in that it leverages outside knowledge to improve performance, and a weakness, in the sense that it makes its application to non-human systems difficult, and may create biases in variant calling.

Here we describe two software programs intended to overcome the limitations of GATK Best Practices, called PEMapper and PECaller. PEMapper solves the inherent limitations of the BWA/GATK pipeline by performing all the necessary read sorting, storing, and mapping procedures in RAM. Human genome indices are preloaded, and final output is written only once (never reloaded, resorted, *etc.*). These technical changes lead to substantial performance gains as detailed below. PEMapper requires a large RAM allocation (typically nearly 200GB for the sequence of a whole human genome), but in exchange, does not over burden the network or disk subsystems. Modern computational clusters, such as those found at many universities or available from cloud providers (i.e. Amazon Web Services), are well equipped to run many simultaneous instances of PEMapper in parallel to expedite experiments. Additionally, output from PEMapper/PECaller comes in much smaller files, decreasing the long-term storage requirements for WGS data. (Table 1)

**Table 1:** Data storage requirements for a single sample using each pipeline

|  |  |  |
| --- | --- | --- |
| **GATK** |  |  |
|  | FASTQ files | 78.9 GB |
|  | BAM file | 115 GB |
|  | Individual VCF file | 53 GB |
|  | Combined VCF file (per sample) | 0.561 GB |
|  | **TOTAL** | **~247.5 GB** |
| **PEMapper/PECaller** |  |  |
|  | FASTQ files | 78.9 GB |
|  | Pileup file | 7.8 GB |
|  | Mapping files | 4 GB |
|  | SNP file (per sample) | 0.035 GB |
|  | Indel file | 0.0001 GB |
|  | **TOTAL** | **~91 GB** |

Unlike PEMapper, whose innovations are strictly in implementation, PECaller represents an intellectual departure from several other genotype-calling models^6,12^ First, variant detection occurs simultaneously (joint-calling in the initial stage) in all samples from the same experiment. This is important because it ensures that the distinction between missing data (data with insufficient evidence for any genotype), and homozygous reference data are recognized from the inception. In addition, it allows the imposition of a population genetics-inspired prior on the data and the ability to fit sophisticated models of read error to help distinguish bases with high error rates from those that actually harbor variants. The population genetics prior accounts for the fact that most sites are expected to be invariant, but conditional on the site containing a variant, the variant is expected to be in Hardy-Weinberg equilibrium. The second innovation of the PECaller method involves the underlying statistical model used to describe the data. Formally, we assume read depths are drawn from a Pólya-Eggenberger (Dirichlet-multinomial) distribution, not the more conventional multinomial assumption. Using a Pólya-Eggenberger distribution allows us to model a nucleotide base both as having a relatively high “error rate,” but also, importantly, a large variance in that rate. This helps us reduce false-positive variant calling, while at the same time enabling us to call true heterozygotes, even when the relative fraction of the two alleles is highly uneven (another common occurrence). We show that PEMapper/PECaller, despite not using any information about “known” SNPs/Indels performs as well or better than GATK Best Practices in all aspects of variant discovery and calling.

## MATERIALS AND METHODS

The PEMapper/PECaller assumes that a reference target sequence is available, but no other information is needed. All mapping and genotype calling occurs relative to this reference sequence. The PEMapper pipeline is composed of a series of three interconnected programs. The first of the three prepares a hashed index of the target sequence. The remaining two programs form a pipeline, with the output of PEMapper forming the input of PECaller. PEMapper is computationally intensive, but extremely gentle on disk and network subsystem. To make this possible, the underlying philosophy behind the PEMapper is that memory usage should be sacrificed for speed and limited I/O. As a result, PEMapper uses approximately 45 bytes of memory per base in the reference sequence, plus approximately one GB of memory per computational core. Therefore, a whole human genome sequence on a 64-core workstation typically uses approximately 200GB of RAM. The source code is freely available at: https://github.com/wingolab-org/pecaller.

The first of the three programs in the PEMapper/PECaller is called *index_target*. Following BLAT^13^, Maq, and several other published algorithms, the target region is decomposed into 16 nucleotide reads. The positions of all overlapping 16-mers in the target are stored. This program needs to be run only once for each target region examined. Unlike GATK Best Practices, this is all that is needed. No information on “known SNPs” or “indels” or training sets is required or used.

The next stage, called *PEMapper*, which also builds on approaches similar to BWA, contains a small innovation to help enable indel mapping. Reminiscent of several other algorithms, the 16-mers are allowed to have up to one sequence mismatch from the target. Thus, when mapping a 100-base read with a 16-base index, an individual read could have up to six errors and still be properly mapped, as long as those errors are evenly distributed along the read. However, the algorithm also allows the 16-mers some “wobble” room, so that relative to each other they can map a few bases away from their expected location (up to eight bases for a 16-mer). Finally, only half of the 16-mers need to map in the correct order, orientation, and distance apart from one another. Positions that satisfy these requirements are taken as “potential mapping” positions.

PEMapper takes this list of putative mapping locations for each read and performs a Smith-Waterman alignment in each potential location to determine the optimal position and alignment score. At this stage, reads are rejected if the final Smith-Waterman alignment score is less than a user-defined percentage of the maximum score possible for the given read length^14^. For results described below, we required 90% of the maximum alignment score and used the following alignment penalties: match = 1, mismatch = −1/3, gap open = −2, and gap extend = −1/36. The primary output of PEMapper is the “pileup” statistic for each base in the target. PEMapper pileup output files include the number of reads where an A, C, G, or T nucleotide was seen, together with the number of times that base appeared deleted, or there was an insertion immediately following the base. Thus, each base appears to have six “channels” of data: the number of A, C, G, T, deletion, and insertion reads.

### The Pólya-Eggenberger distribution

The Pólya-Eggenberger (PE) distribution is a multidimensional extension of the beta-binomial distribution. Although it arises in numerous contexts and was initially described in connection with an urn sampling model^15^, for our purposes we view the PE distribution as the result of multinomial sampling when the underlying multinomial coefficients are themselves drawn from a Dirichlet distribution^16^, in the same way the one-dimensional analog, the beta-binomial distribution, can be thought of as binomial sampling with beta-distributed probability of success. Intuitively, we envision the six channels of data (number of A, C, G, T, deletion, and insertion reads) as being multinomially sampled with some probability of drawing a read from each of the channels, but that the probability varies from experiment to experiment and is itself drawn from a Dirichlet distribution. The coupling of the Dirchlet distribution with the multinomial distribution is common in Bayesian inference, as the former distribution is often used as a conjugate prior for parameters modeled in the latter distribution^16^. Here, our purpose is subtly different. In Bayesian estimation, the assumption is that the observations are fundamentally multinomial, but the parameters of that multinomial are unknown, and the Dirichlet is used to measure the degree of that uncertainty in the parameter estimates. In the Bayesian estimation case, as the data size gets sufficiently large, convergence to a multinomial occurs. Here, on the other hand, we assume that the observations are fundamentally over-dispersed relative to a multinomial, and there is not necessarily a multinomial convergence.

At any given base, a diploid sample could be one of 21 possible genotypes (a homozygote of A, C, G, T, deletion, or insertion, and all 15 possible heterozygotes). We assume that the number of reads seen in each of the six possible channels (A, C, G, T, Deletion, Insertion) of data for an individual with genotype *j* is drawn from a PE distribution in six dimensions. We further assume that each of the 21 possible genotypes is characterized by its own PE distribution, and that these 21 distributions vary from base to base, but are shared by all samples at a given base. A six-dimensional PE distribution is characterized by six parameters, so let **a_*j*_** be a six-dimensional vector corresponding to the parameters for genotype *j*. If **n_*i*_** is a six-dimensional vector containing the six channels of data observed in individual *i* at a given base, and if individual *i* has genotype *j*, then the probability of those observations is

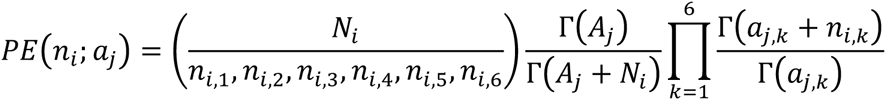

where *N_i_* is the total number of reads observed (*N_i_* = *n_i,1_+n_i,2_+n_i,3_+n_i,4_+n_i,5_+n_i,6_)* for individual *i*, *A_j_* is the corresponding sum of the parameters for genotype *j* (*A_j_*=*a_j,1_+a_j,2_+a_i,3_+a_j,4_+a_j,5_+a_j,6_)*, and Γ, is the usual gamma function^17^. Note that the expected proportion of reads coming from channel *k* is given simply by *a_j,k_* /*A_j_*.

### Genotype calling overview

Genotype calling occurs across all samples simultaneously in a fundamentally Bayesian, but iterative, manner. First, the PE parameters for all 21 genotypes are set to “default values” and assumed to be known. Second, the genotypes of all the samples are called in a Bayesian manner, conditional on the “known” PE parameters. Finally, the PE parameters are estimated, conditional on the genotypes called in step two. The process then iterates, with the genotypes re-called, and parameters re-estimated. The iteration continues until either calls no longer change, or a maximum number of iterations is reached. For all the results described here, the maximum was set at five iterations, which was seldom reached.

### PE parameter initialization

For all 21 genotype models, *A_j_* is set to either the average read depth across samples or 100, whichever is larger. For homozygote base calls (A, C, G, T, but not indels), the expected proportion of reads coming from channels different from the channel associated with the homozygote allele (*i.e.*, the expected proportion of error reads) is set at 1/*A_j_* or 0.3%, whichever is larger for each channel; thus, at initialization we assume between 0.3% and 1% “error” reads in every channel. The remainder of the reads are expected to come from the “correct” channel. For heterozygote genotype calls, the error channels are set similarly, except for the “deletion” channel, which is expected to have 5% of the reads, indicating a prior assumption that approximately 5% of true heterozygous reads will map incorrectly as deletions. If the heterozygote genotype does not involve the reference allele, the remaining reads are expected to come equally to both of the appropriate channels. On the other hand, if the heterozygote includes the reference allele, we assume that 52% of the remaining reads map to the reference allele, and 48% to the non-reference. This incorporates our notion that some portion of the time non-reference alleles will not map, or map incorrectly as indels.

To meet the challenge of mapping indel variation, we made the following assumptions: for deletion homozygotes, we again assume a 0.3%-1% read proportion in all the channels that do not involve the reference allele or the deletion; however, we expect the remaining reads to divide 80% deletion and 20% reference, indicating our assumption that a substantial fraction of deletion reads mis-map as reference, even when the deletion is homozygous. When the deletion is heterozygous, we assume the non-error channels to divide 60%-40% between the reference channel and the deletion channel. Insertions after the current base are again assumed to have 0.3%-1% reads in the error channels. For homozygotes, 80% of the remaining reads are expected to include the reference base and have an insertion afterwards, whereas 20% of the reads will only include the reference allele. For heterozygous insertions, 40% of the remaining reads are expected to include both the reference allele and an insertion, and 60% only the reference allele.

### Bayesian genotype calling with a population genetics prior

We assume that *m* samples have been sequenced. Each of those *m* samples can be any of 21 possible genotypes. Thus, there are a total of *(21)*^*m*^ possible genotype configurations of those *m* samples. Let **c_k_** be one such configuration. **c_k_** is an *m*-dimensional vector, where element *c_k,i_* is an integer between 1 and 21, and indicates the genotype of sample *i*. Genotypes of all the samples are assumed to be independent, and therefore the likelihood of configuration **c_k_** is

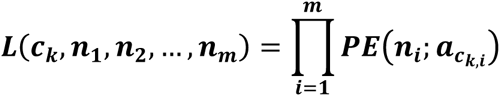

Most sites will not be segregating, and all *m* samples will be identical to the reference allele. Let **c_0_** be the configuration where all samples are the reference allele. By assumption, the prior probability that this configuration is

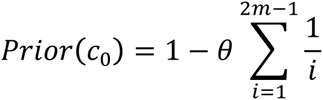

where θ is a user-supplied parameter corresponding to 4N_e_μ, N_e_ is the effective size of the population from which the samples were drawn, and μ is the per-site per-generation mutation rate^18^. For humans it is generally assumed to be ~0.001^19^. All other configurations have at least one sample with at least one allele different from the reference allele. Let f(**c_k_**,r) be the number of non-reference alleles of type *r*, *0 < r < 6*, found in configuration **c_k_**. The prior probability of configuration **c_k_** is assumed to be

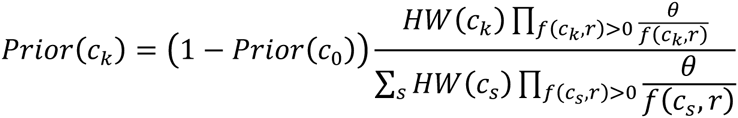

where HW(**c_k_**) is the Hardy-Weinberg exact p-value^20^ associated with configuration **c_k_**, and the sum in the denominator is taken over all *(21)*^*m*^ - 1 genotype configurations (but see computational efficiencies section below).

Overall, this prior can be summarized as follows. The population from which these samples are drawn is assumed to be of constant size and neutral, and the reference allele is assumed to be the ancestral allele at every site. The prior probability that a site is segregating is the one derived by Watterson for an infinite-site neutral model^18^. Conditional on the site segregating, the assumption is that the site is in Hardy-Weinberg equilibrium, and the derived allele frequency was drawn from an infinite-site neutral model. Thus, the prior probability is a combination of two terms, one of which derives from the Hardy-Weinberg p-value, and the other from the number of different alleles seen to be segregating. Finally, we should note that we have tacitly assumed that all the sequenced samples are randomly drawn from the underlying population, *i.e.*, not intentionally picked to be relatives of one another. Alternatively, the user may provide a standard linkage/ped (PLINK pedigree format)^21^ to specify the relationship between samples. When this option is invoked, Hardy-Weinberg is calculated only among unrelated individuals (*i.e.* founders), and for every configuration, **c_k_**, the minimum number of *de novo* mutations, **Dn(c_k_)**, is calculated for the configuration. Each *de novo* mutation is assumed to occur with user specified probability μ, and the prior probability of the configuration is modified to

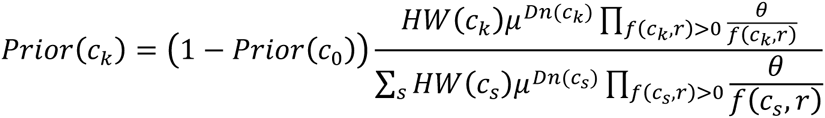

The posterior probability of configuration **c_k_** is

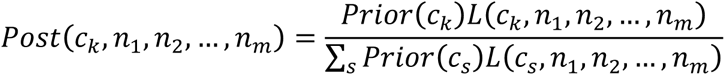

where the sum is taken over all possible genotype configurations (but see below). If 0 < *g_i_* < 22 is the genotype of individual *i*, then

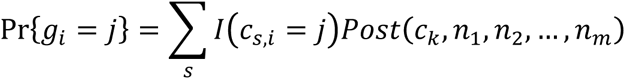

where *I(c_s,i_=j)* is an indicator function that equals 1 whenever element *i* of configuration **c_s_** is equal to *j*, and is 0 otherwise. Thus, we take the probability that the genotype of individual *i* is *j* to be the sum of the posterior probabilities of the genotype configurations in which we call sample *i* genotype *j*. The PECaller calls sample *i* genotype *j* whenever *Pr(g_i_=j)* is greater than some user-defined threshold, and otherwise the genotype is called “N” for undetermined. For all the results presented here, the threshold was set at 0.95.

### Estimating PE parameters and repeating

Because of local sequence context, the repetitive nature of many organisms’ sequence, and specific issues with sequencing chemistry as a function of base composition, not all bases have the same “error” characteristics. Some bases may appear to have a very high fraction of reads containing “errors,” while other bases have almost none. Some heterozygotes may exhibit nearly 50-50 ratios of the two alleles; others can be highly asymmetrical. To account for this, we wish to estimate the PE parameters independently at every base. There are three technical challenges to this. First, and most importantly, the genotypes of the samples are not known with certainty, hence we do not know with certainty which observations are associated with which underlying PE distribution. Second, for technical reasons (one lane “worked better” than another, *etc.*) some samples may have many more reads than other samples, and we do not want these high-read samples to dominate our estimates disproportionately. Finally, because it is necessary to estimate parameters repeatedly, the algorithm must be computationally efficient. With this in mind, we chose moment-based estimators of our parameters^22^.

In principle, we would like to estimate the PE coefficients for genotype *j*, **a_*j*_**, by averaging over the observed number of reads seen in every sample that has genotype *j*; however, we do not know this with certainty. So, let **f_i_** be a six-dimensional vector, where element f_*i,k*_ = *n*_*i,k*_/*N*_*i*_ contains the fraction of individual *i*’s reads that were observed in channel *k*. Let

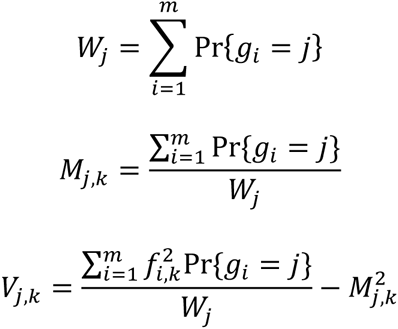

Thus, *M_j,k_* and *V_j,k_* are the “weighted” mean and variance in read fraction from channel *k* among individuals with genotype *j*, where both moments are “weighted” by our confidence that the individual truly is genotype *j*. Usually, most genotypes will have little weight (*i.e.*, few if any samples are called that genotype), and even when samples are called that genotype, sometimes there is little to no variation seen in read fractions (*i.e.*, 100% of the reads come from one channel in all the samples called that genotype). Let *Y_j_* be the number of channels for genotype *j* that have non-zero observed variance in read fraction. Thus,

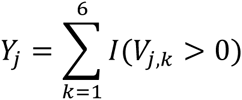

where *I(V_j,k_ > 0)* is an indicator that genotype *j* has non-zero variance in channel *k.* For any genotype with *W_j_ < 1.5* (*i.e.,* less than two samples called that genotype), or with *Y_j_* < 2 (*i.e.*, less than two channels with variance in read fraction), all PE parameters are returned to their initialization values. Otherwise, let channel *z* be the channel with non-zero variance (*V_j,z_* >0), but minimal mean (*M_j,z_* < *M_j,k_*, for all other *k* with non-zero variance) estimate

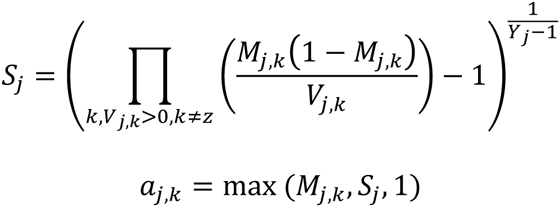

S_j_ can be thought of as a “leave one out” moment estimate of the “precision” of the PE distribution, and *M_j,k_* is a first-moment estimate of the mean read fraction in each channel^22^. Notice that all channels with a small expected read fraction are rounded up to one (see below). Once the PE parameters for all the genotype models are estimated, the process repeats, and genotypes are re-called, until genotype calls no longer change, or a maximum of five iterations is reached.

### Computational efficiencies

The sample space of configurations is impossibly large. For anything other than a trivially small number of samples, the sums over the configuration sample space cannot be done. Nevertheless, the prior distribution is remarkably “flat,” and this can be used to great advantage. If two configurations, **c_u_** and **c_v,_** differ by only a single sample’s genotype, then we know that the ratio of their prior probabilities is bounded by

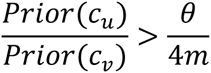

To see this note that the largest difference in prior probabilities occurs when configuration **c_u_** has a single homozygote of an allele not seen in configuration **c_v_**. The difference in Hardy-Weinberg p-values associated with this is less than 1/2m^20^, and the difference due to the number of alleles segregating is θ/2. Thus, if

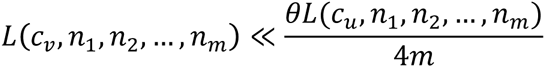

then

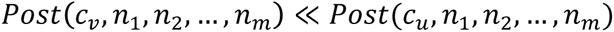

The immediate implication of this is that dropping configuration **c_v_** from the sum will have little effect on the posterior probabilities of any of the *likely* configurations of the genotypes, and a simple, nearly linear time algorithm to enumerate all the likely configurations and ignore the unlikely ones is suggested.

We build the list of likely configurations by moving through the samples one at time. Initially, we start with a set of 21 configurations that correspond to all the possible genotypes for sample 1. We calculate the likelihood of all 21 one-sample configurations, and then remove any configuration with likelihood less than 10^−6^ times the largest likelihood. Additionally, we always save the configuration associated with all samples being homozygote reference, because *a priori* this is the most likely configuration of samples. Next, to each of the remaining configurations we add all 21 possible genotypes for the second sample, thereby increasing the number of sample configurations by a factor of 21. However, we again immediately remove all configurations with likelihood less than 10^−6^ times the largest likelihood. We repeat until we have gone through all *m* samples. In principle each step could increase the number of likely configurations by a factor of 21, but in practice it almost never increases the number by more than a factor of two (*i.e.*, there are almost never more than two likely genotypes for one sample), and most of the time it does not increase the number of configurations at all (*i.e.*, most of the time there is only one likely genotype for a sample). Even when *m* is in the hundreds, most bases have only a handful of likely configurations, and seldom is the total number of likely configurations more than a few thousand.

PECaller takes advantage of two other computational efficiency tricks. First, HW exact probabilities are fundamentally discrete and a simple function of the number of heterozygous and homozygous genotypes. Those values can be calculated ahead of time and stored in lookup tables, greatly aiding that computation. Second, Pólya-Eggenberger distributions contain several gamma functions, and although gamma functions can be computationally expensive to calculate, in a special case, they are cheap. If *x* is an integer, Γ(x) is equal to *(x-1)* factorial, so we round all PE coefficients to their nearest integer greater than or equal to one. PE distributions can be calculated strictly in terms of factorials, and it is easy to precalculate and store all factorial values less than, say, 10,000. It should be noted, as well, that all likelihood calculations occur computationally as natural logs and are raised to an exponential only when necessary for posterior probability determinations. Thus, as a practical matter, the natural log of factorials is computed and stored.

Finally, both PEMapper and PECaller can be set to disregard highly repetitive sequences. By default, during the initial placement of reads, PEMapper ignores any 16-mer that maps to over 100 different locations in the genome. Thus, in order to even attempt Smith-Waterman alignment, at least half of the 16-mers in a read must map to less than 100 places in the genome. Any read more repetitive than this is dropped. Similarly PECaller can be given a file in bed format that constitutes the “target” region to be called. This can be used, for example, to specify the exome-only, for exome studies, or the non-repeat masked regions of the human genome for WGS studies. Since variation in repeat-masked regions is both extremely difficult to interpret, and highly prone to error/mismapping, all the results describe will be for the unique portion of the genome (i.e. non-repeat-masked).

### Bisulfite sequencing and other user options

A possible application of next-generation sequencing is to determine the pattern of methylation in a given region sequenced. One way of doing this is to first treat the DNA with bisulfite, which converts C’s to T’s, unless the C has been methylated. Bisulfite treatment can pose unique challenges for mapping short sequence reads. The PEMapper/PECaller contains a user option to gracefully handle bisulfite-treated DNA. When the user selects this option, all mapping is initially done in a “three-base genome,” where C’s and T’s are treated as if they are the same nucleotide. Indexing of the genome is done in this three-base system, as is initial mapping. Final placement of reads with Smith-Waterman alignment is done in a four-base system, but C-T mismatches are scored as if they are perfect matches. The methylation status of any C allele can then be immediately calculated from the “pileup” files, which gives the number of C and T alleles mapping at any base.

Many second-generation sequencing technologies can create both single-ended and pair-ended reads, with either single files per sample, or multiple files per sample. The PEMapper can take all these forms of data, and for pair-ended data, the user specifies the minimum and maximum expected distance between the mate-pair reads. For mate-pair data, the PEMapper will first attempt to place the reads in a manner consistent with the library construction rules, but if no such placement can be made, it will place one or both reads if they individually map uniquely with sufficiently high score.

Throughout the genotype calling section, we assumed that every sample was diploid, and therefore that there were 21 possible genotypes for any sample at a given base. If the user specifies that this is haploid data, only six possible genotypes are assumed (homozygotes for any of the six alleles), and the Hardy-Weinberg p-value is removed from the prior.

### WGS

We tested the performance of GATK and PEMapper on 97 WGS samples, sequenced as part of the International Consortium on Brain and Behavior in 22q11.2 Deletion Syndrome (IBBC; www.22q11-ibbc.org). The collaboration, an initiative supported by the National Institute of Mental Health, combines genomic with neuropsychiatric and neurobehavioral paradigms to advance the understanding of the pathogenesis of schizophrenia (SZ) and related disorders, given the high risk for these conditions (> 1 in 4), in individuals with the 22q11.2 deletion^23^. Rigorous approaches are applied across the IBBC to characterize the phenotypes, the 22q11.2 deletion and the remaining genome. DNA samples from 97 participants ^24,25^ each with a typical 2.5 Mb hemizygous 22q11.2 deletion. Eight of these participants have previously published WGS data using different methods^25^.

All samples were sequenced at the Hudson-Alpha Institute of Biotechnology (HAIB, Birmingham, AL) on Illumina HiSeq-2500 machines, using their published protocols. Briefly, the concentration of each DNA sample was measured by fluorometric means (typically PicoGreen reagent from Invitrogen), followed by agarose gel electrophoresis to verify the integrity of DNA. Following sample quality control, all samples with passing metrics were processed to create a sequencing library. For each sample, 2 µg of blood-extracted genomic DNA was sheared with a Covaris sonicator, the fragmented DNA was purified, and paired-end libraries were generated using standard reagents. Yields were monitored following sonication, ligation, and at the complete library stage with additional PicoGreen quantitation steps. Every library in the project was tagged with a two-dimensional barcode that leverages the Illumina sequencer’s ability to perform four sequencing reads per run (two data reads and two index reads). Two types of quality control were performed on each library prior to sequencing. First, the size distribution of the library was determined with a Perkin-Elmer/Caliper LabChip GX to verify a correctly formed and appropriately sized library. To avoid overlapping reads, a physical size of 500-600 bp was verified on the Caliper or Agilent instrument. This observed physical size corresponds to an alignment-based insert size of slightly over 300 bp. The second step in the quality-control process was a real-time, quantitative PCR assay with universal primers to precisely quantity the fragments that are able to be sequenced in the library. The real-time PCR results, in combination with the size data, were used to normalize all libraries to a 10-nM final concentration. Following quality control, each plate of 96 libraries was pooled into a single, complex pool. The final library pool was sequenced on a test run using the Illumina MiSeq instrument and a paired-end 150-nt sequencing condition with indexing reads. The data from the MiSeq served as a final quality control step for both samples and the libraries. Libraries that passed QC were subjected to full sequencing on the Illumina HiSeq 2500 instruments according to current Illumina protocols, essentially as described in Bentley, 2008^26^. The unique barcoding features of the described library construction allow up to 96 samples to be pooled and sequenced simultaneously. Of these samples, 93 were also run on Illumina Omni 2.5 genotyping arrays (http://www.illumina.com/techniques/popular-applications/genotyping.html) which served as an additional sequencing quality control.

### PEMapper/PECaller methods

PEMapper was run on Amazon Web Services r3.8xlarge instances with 32 CPUs with 244GB of RAM for each sample. Globus Genomics (www.globus.org) was contracted to facilitate the running of PEMapper on AWS. A PEMapper workflow is available through Globus, which leverages batch submission, such that multiple samples can be submitted for mapping simultaneously. The sequencing files (fastq format) were uploaded to AWS via Globus, and the PEMapper output is subsequently returned to the user’s local machine. PEMapper was run with all default parameters, and a 90% threshold for Smith-Waterman alignment. PECaller was run with a default theta value of 0.001 (See results), and a 95% posterior probability for a genotype to be considered called (less than 95% is reported as “missing” or “N”). Sites with less than 90% complete data were dropped. All mapping and genotyping occurred relative to the human HG38 reference, as reported by the UCSC Genome Browser on July 1, 2015. We report results only for the non-repeat-masked portion of the genome.

### End user instructions

Running the PEMapper/PECaller pipeline is very straightforward for an end user. One begins with fastq files from whole-genome sequencing (the number doesn’t matter, how ever many represent the complete sequencing of the sample of interest). If the end user has opted to use the Globus Genomics pipeline on AWS, the fastq files are uploaded to the PEMapper workflow and the user receives three important files in return: a pileup file, a summary file, and an indel file. If the end user is running PEMapper locally, he or she must have a copy of the reference genome and load that into memory before running PEMapper with the map_directory_array.pl script. In either case, the user will run PECaller locally. To do so, one gathers the pileup and indel files for each sample to be processed in a single folder. The script call_directory.pl is used to launch PECaller. That generates a .snp file (containing all SNPs, but no indels, in an unsorted list) as output. Then merge_indel_snp.pl is run to merge the indels into the list of SNPs. This produces a merged snp file (containing SNPs and indels in a sorted list). This file can be converted simply to a PLINK pedigree format, and represents the primary output of the pipeline. Several additional scripts permit easy quality control assessments of the data. The first script, snp_tran_counter.pl generates a file with transition-to-transversion (Ts/Tv) information about the samples. At this point, the web-based annotation program, Seqant (http://seqant.genetics.emory.edu/)^27^ can be used to annotate the merged .snp file. Finally, a second script, snp_tran_silent_rep.pl takes the output from SeqAnt and generates a file with silent/replacement information about the samples.

### GATK methods

The initial steps of GATK, BWA and Haplotype Caller, were similarly run on Amazon Web Services r3.8xlarge instances with 32 CPUs with 244GB of RAM for each sample. Globus Genomics (www.globus.org) was also contracted to facilitate the running of GATK. A GATK workflow is available through them that runs, in order, BWA v0.7.12-r1039, sambaba v0.5.4, and GATK v3.5-0-g36282e4. The reference genome utilized was hg38, downloaded from the Broad Institute. This workflow leverages batch submission, such that multiple samples can be submitted for mapping simultaneously. The sequencing files (fastq format) were uploaded to AWS through Globus and the GATK output (BAM and VCF files) was subsequently returned to the user’s local machine.

Joint genotyping and variant recalibration were done in GATK v3.6 locally, in batches of 10 samples due to the intensive computational load. The joint genotyping and variant recalibration tools were run on nodes with 64 cores and 512GB of RAM. All mapping and genotype calling was relative to same reference hg38 genome in PEMapper/PECaller, with SNP sets, etc taken from the hg38 resource bundle provided by GATK. All repeat-masked regions of the genome were dropped. The Unified caller would not run on the entire 97 sample dataset even on compute nodes with 512GB of RAM free (it always eventually reported an “out of heap space” error whether run on the whole genome or each chromosome separately). We attempted to run the unified caller on subsequently smaller batches of data: it would complete in a batch size of 10 genomes, but failed at a batch size of 20. Results below are from nine batches of 10 samples each, and one batch of seven.

## RESULTS

### Performance of PEMapper/PECaller

The simplest measure of variation, named theta^18^, counts the number of heterozygotes called per sample per base. Theta is estimated to be somewhere between 0.0008 in 0.001 in humans^19,28^. Figure 1 shows theta for each of the 97 sequenced human genome samples that passed QC (see Methods). Most remarkable is the extremely consistent levels of variation called between samples, with individuals ranging from .0007899 to .0008204. The overall variation levels are consistent with previous estimates.

**Figure 1:**
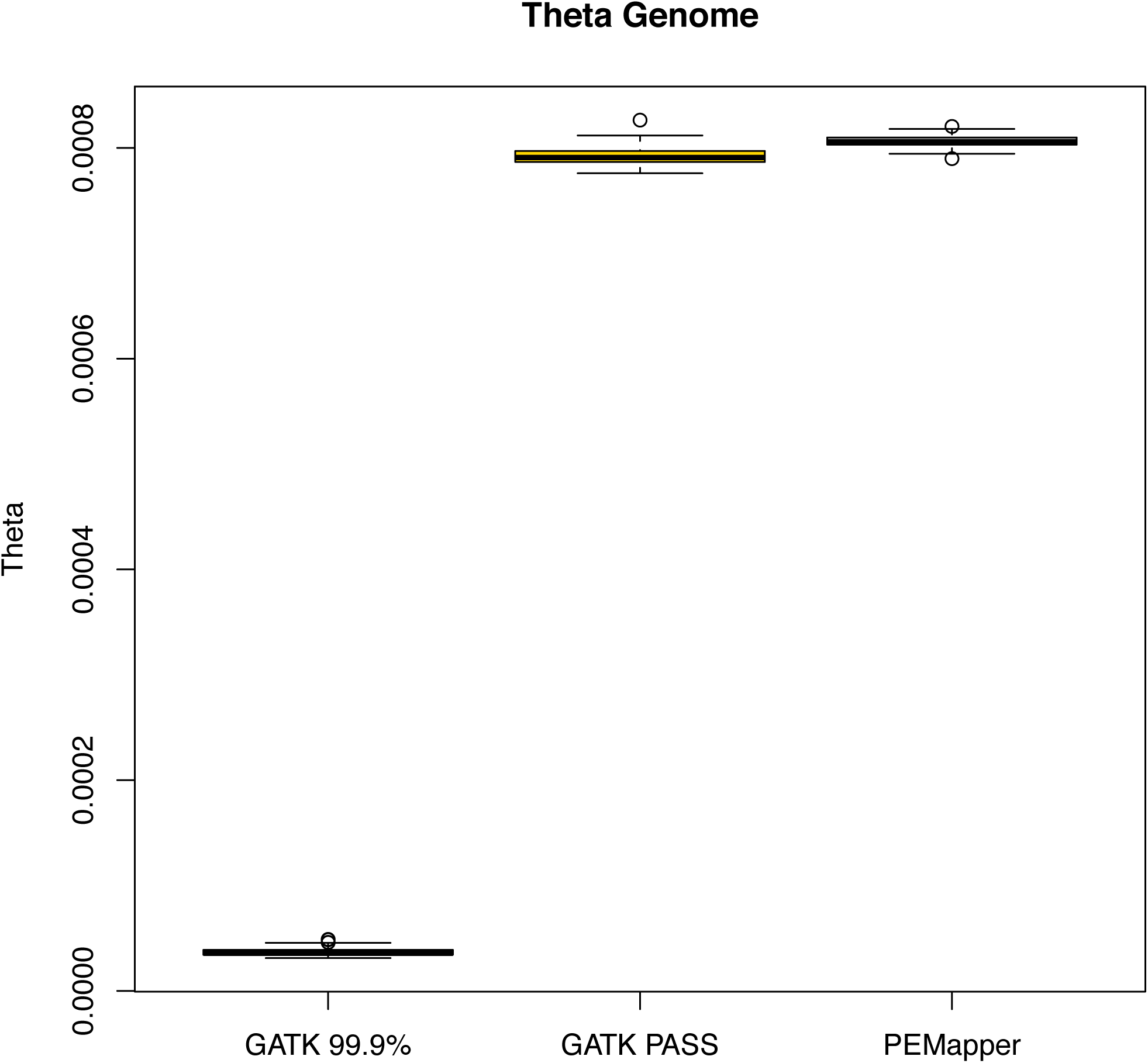
Theta across all samples. Theta across all 97 samples based on the calls from PEMapper/PECaller, GATK PASS and GATK Tranche99.9. PEMapper/PECaller and GATK PASS samples sit between .00075 and .0009 variants per base, as expected. Tranche99.9 calls are much lower.

### False-positive calling

Our analysis provides ample evidence that this called variation contains very few false positive findings (non-variant sites called variant in error). Sequence changes from A->G, G->A, C->T, and T->C are called “transitions.” All other changes are called “transversions.” There are twice as many transversions possible as transitions. Many mutational mechanisms favor transitions over transversions (oxidative deamination, etc.). Selection also likely favors transitions over transversions (much more likely to be silent in exons, more similar binding for transcription factors, e.g. wobble binding). On the other hand, random genotype calling error likely results in increased transversions (because there are twice as many ways to get a transversion as a transition when you make an error). Thus, real data ought to be enriched for transitions over transversions, and false data ought to be enriched for transversions. Picking nucleotides at random would give a 0.5:1 transition-to-transversion ratio. It is widely believed that the overall transition-to-transversion ratio is approximately 2.0 in humans (http://genome.sph.umich.edu/wiki/SNP_Call_Set_Properties). For every sample in this study, the transition-to-transversion ratio was between 2.042:1 and 2.051:1 (Figure 2). Looking at the entire collection of variants, the ratio was 2.073:1. This overall ratio can be used to estimate the fraction of false-positive variant calls. If we assume the “true” ratio is 2.12:1, a value determined from all variants called both by PEMapper/PECaller and GATK (see below), and we assume that false-positive variant calls have a ratio of 0.5:1 (as expected by chance), then an observed ratio of 2.073:1 implies that, over the entire 97 samples, approximately 3% of the variants were false positives. On a per-sample basis, less than 1 in 3000-5000 called variants per sample were false positives. The data quality from PEMapper/PECaller compares favorably to other NGS analytical tools^29^.

**Figure 2:**
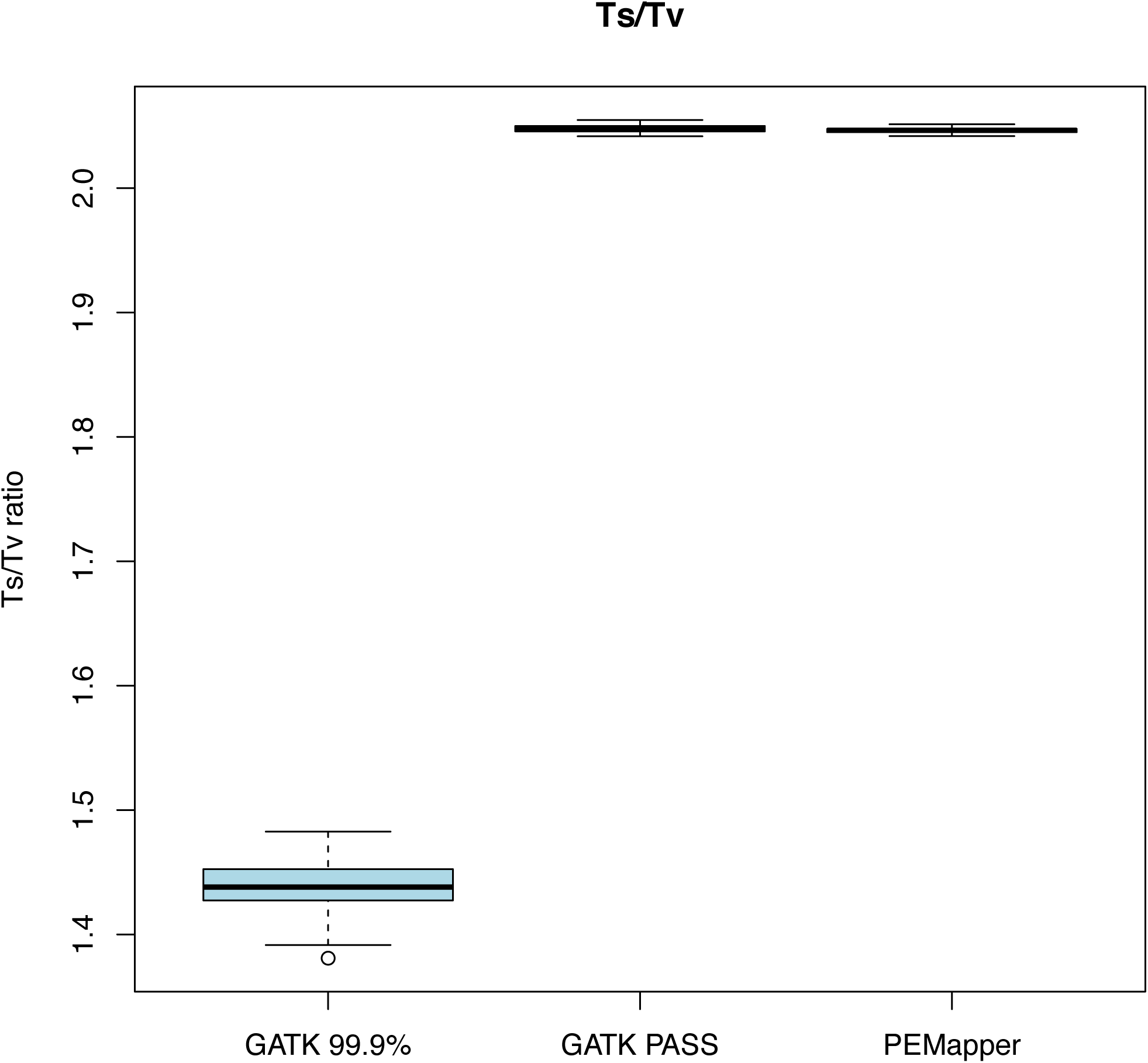
Comparison of Ts/Tv ratio across all calling results. Comparison of transition-to-transversion ratios for PEMapper/Caller, GATK PASS and GATK Tranche99.9 called variants. PEMapper/PECaller and GATK PASS are virtually identical at near 2.04 and 2.05 per sample, indicating excellent quality calls. GATK Tranche99.9 is much lower, between 1.3 and 1.5 per sample, indicating much lower quality calls.

### Exonic variation

In general there ought to be far less variation in exons than in the genome as a whole. In these samples, we saw Theta in exons to be between 0.0004284–0.0004550 per sample (Figure 3), i.e. slightly more than half its value for the genome as a whole. We also found a much higher transition-to-transversion ratio (2.963:1 to 3.130:1) (Figure 4), consistent with selection for transitions in exons. Of the variants in exons, one expects approximately half to be “silent” (making no change to the amino acid sequence) and half to be replacement (changing the amino acid sequence). The average silent-to-replacement ratio^27^ per sample was 1.101:1, with a range from 1.074:1 to 1.127:1 (Figure 5). On average, there were approximately ~20,000 variants in the CCDS-defined (Consensus Coding Sequence Project, https://www.ncbi.nlm.nih.gov/projects/CCDS/CcdsBrowse.cgi) exome of each individual. Over the entire collection of sites, 44.54% of all exonic variants were silent. This number is remarkably similar to published estimates from 100x whole-exome sequencing^30^. Of note, Tennessen et al. restricted themselves to ~16,000 well-covered genes, where here we use the whole CCDS exome.

**Figure 3:**
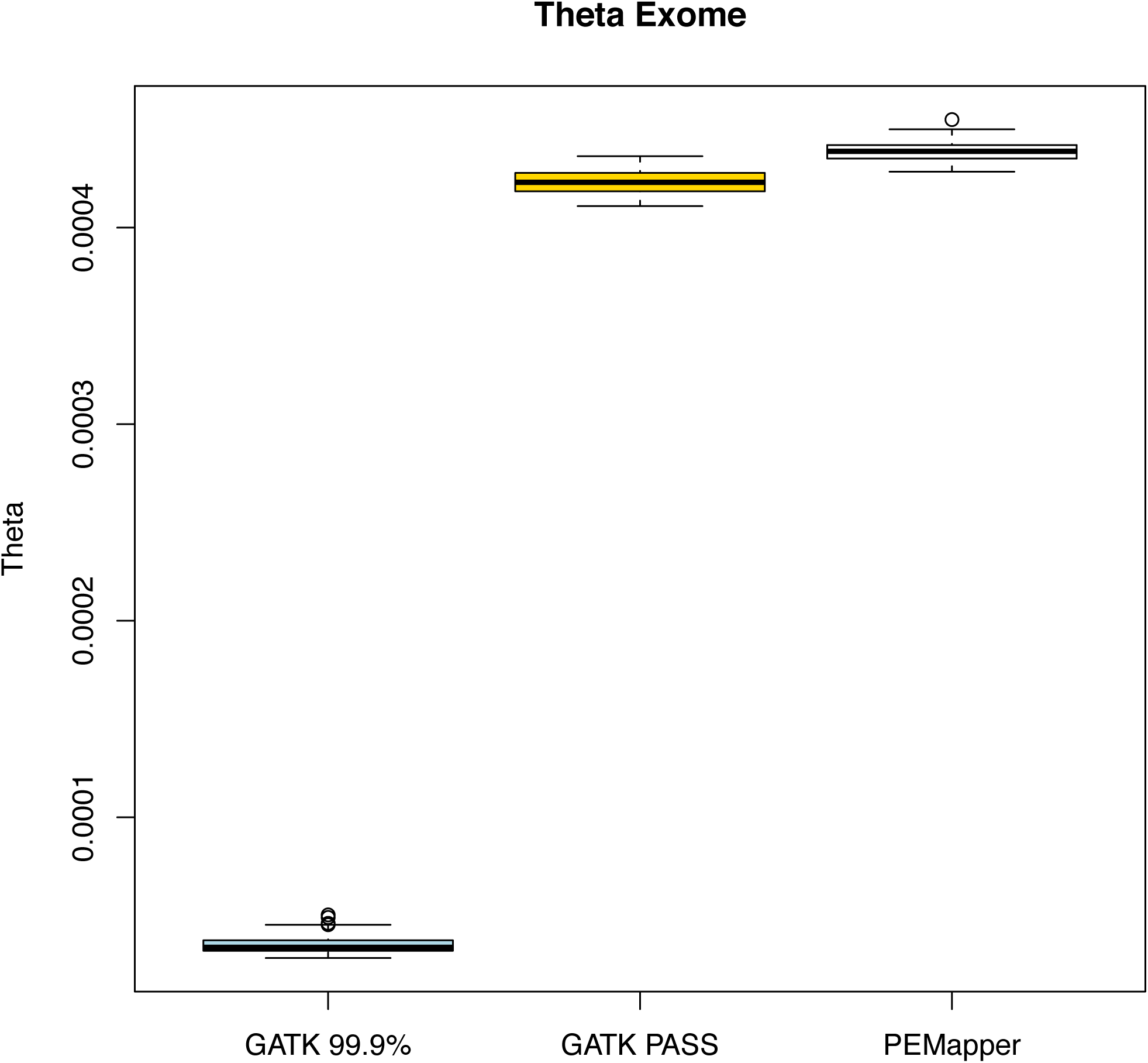
Exomic theta. Theta in all sample exomes based on PEMapper/PECaller, GATK PASS and Tranche99.9 calls. GATK PASS and PEMapper/PECaller samples are near .00045, as expected, with PEMapper/PECaller calling slightly more variants.

**Figure 4:**
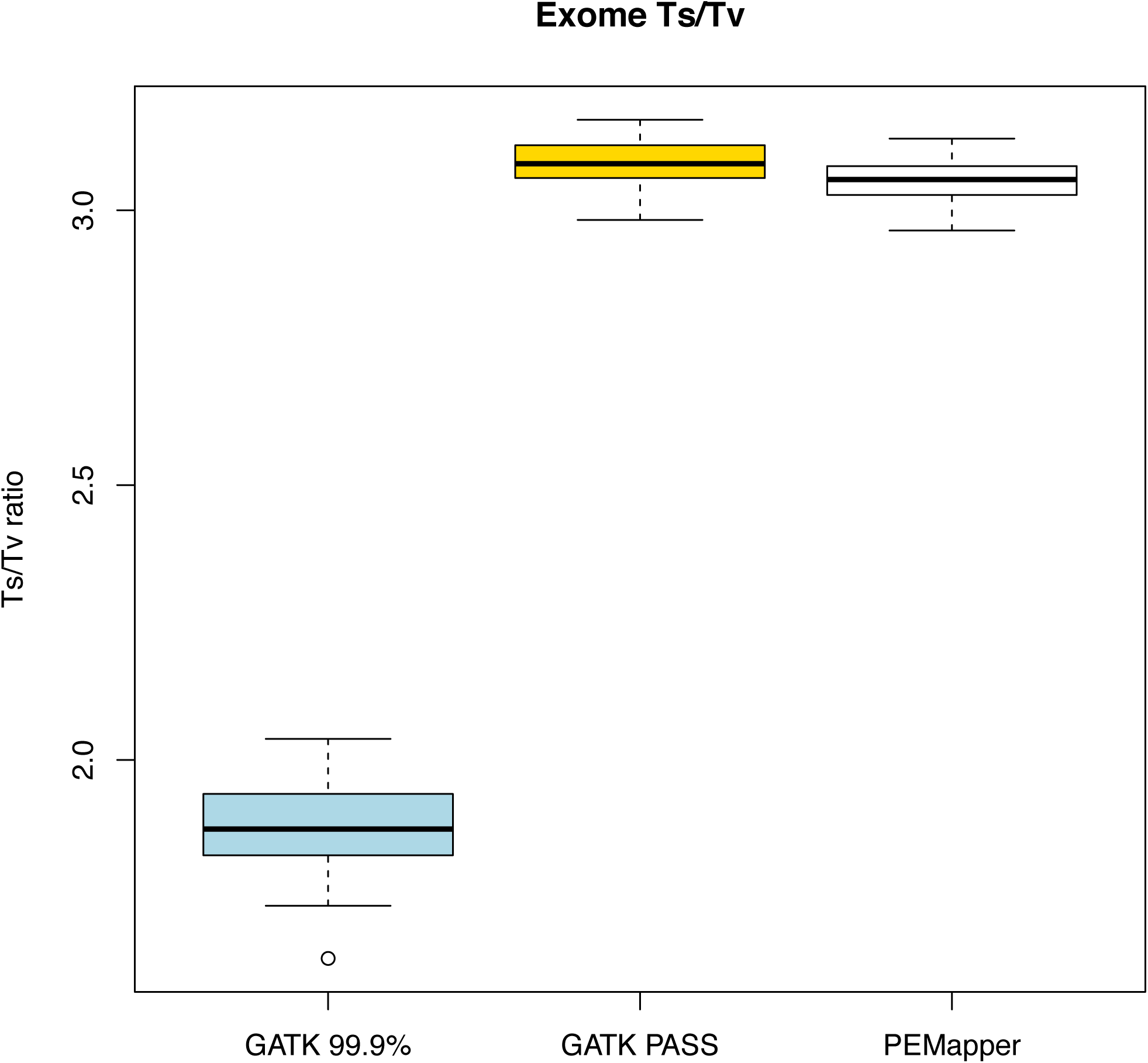
Exomic transition-to-transversion ratio. Transition-to-transversion ratio across all sample exomes based on PEMapper/PECaller, GATK PASS and Tranche99.9 calls. All samples called by PEMapper/PECaller and GATK PASS are near 3, as expected. Tranche99.9 calls are much lower, again.

**Figure 5:**
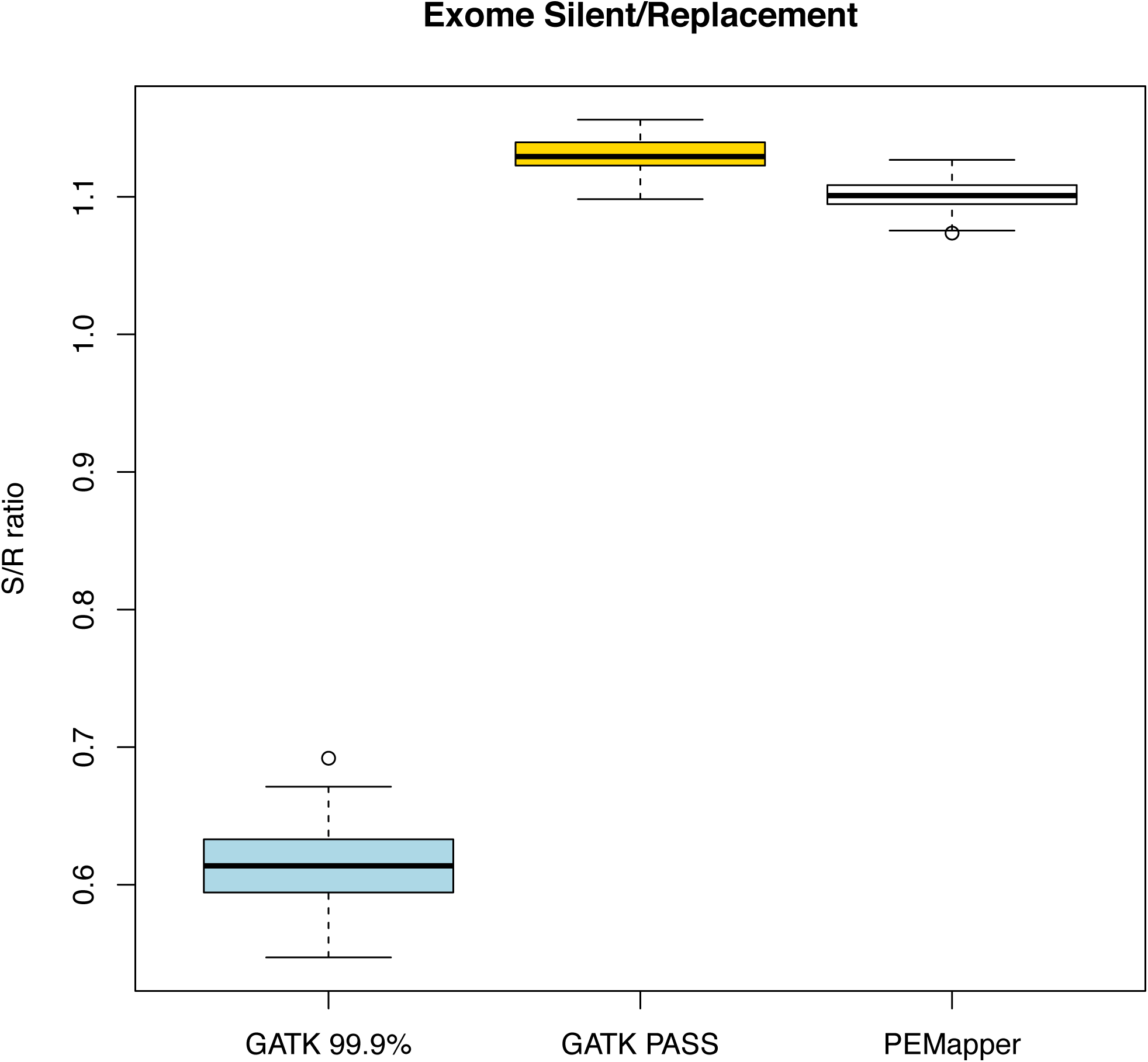
Exomic silent-to-replacement ratio. Silent-to-replacement ratio across all sample exomes based on PEMapper/PECaller, GATK PASS and Tranche99.9 calls. All samples called by PEMapper/PECaller and GATK PASS are between 1.05 and 1.15, as expected. Again, Tranche99.9 calls are significantly lower.

### Calling Rare Variation

Naively we might imagine most false positives to be in the “singleton” category, *i.e.* variants seen only once in our sample set. Here, singletons have a Ts/Tv ratio of 2.105 to 1, better than the PEMapper/PECaller average of 2.073 to 1, and very close to Ts/Tv ratio of the overlap set between GATK and PEMapper/PECaller. So singletons, despite the additional potential for false positive calls, actually appear to be as reliable or more reliable than the set of all sites.

dbSNP 146 contains all variants currently reported in the ExAC^7^ dataset, as well as all variants discovered by 1000 Genomes^31^. An exonic variant not found in dbSNP 146 is almost surely either a false positive call, or a variant that is exceedingly rare in the general population. Exonic sites that change the amino acid (replacement sites) and are not found in dbSNP should be the category of variation most enriched for false positive calls. For the entire set of replacement SNPs, the Ts/Tv ratio is 2.173. Replacement SNPs in dbSNP are 2.254, while those not in dbSNP are 1.762. For singleton replacement SNPs, the Ts/Tv ratio is 2.328. Singleton, replacement SNPs in dbSNP are 2.562, while those not in dbSNP are 1.846. This set of singleton replacement sites that are not found in dbSNP is the set that ought to be most enriched for false positives. In spite of this, replacement sites that are not in dbSNP have a TS/TV ratio only ~10% lower than SNPs overall, suggestive that while this set may be the most enriched for false positives of any possible set, it is still comprised largely of true positive calls.

### Completeness and accuracy

Overall, more than 98.4% of the non-repeat-masked genome had high-quality calls. As expected, more than 99% of these sites were called homozygous reference in all 97 samples. At sites called variant in at least one sample, our overall data completeness was 99%. Most of these samples (93) were also genotyped on Illumina 2.5M arrays. These arrays provide over 140 million genotypes that can be compared to the sequence-called genotype. Over these 140 million genotype calls, PECaller data were 99.85% complete, and agreed with array call 99.76% of the time. Partitioning these numbers by array-called genotype, we note that if the genotyping array called “homozygote reference,” the sequencing call was 99.95% complete and agreed 99.94% of the time. If the array called a “heterozygote,” the sequencing was 99.81% complete and agreed 99.23% of the time. Finally, if the array called a “homozygote non-reference,” the sequencing was 99.88% complete and agreed with the array 99.56% of the time.

Lack of agreement between sequencing- and array-based calls can be due to errors in either the array or the sequencing call. One can show that, if the arrays are 99.8% accurate regardless of true genotype, the agreement level above is consistent with sequencing being 99.9% accurate overall, i.e. if arrays are only 99.8% accurate, most of the disagreements between array and sequencing are due to array errors.

### Rare variant false negatives

While the overall completeness and accuracy at high-frequency sites is excellent (99.85% complete and more than 99.76% accurate), it is possible that data completeness and accuracy at low-frequency variants might be considerably worse. This could occur because joint calling of samples can increase one’s confidence for high-frequency variants, while providing comparatively little benefit for rare variant calling. To assess the probability of “missing” rare variants, we look at variants called by the Illumina 2.5M array where the variant allele was observed in only one of our samples. In this collection of ~40,000 “singleton” variants, we do not see evidence for increased missing data rates in singleton variants, with only 0.24% missing data. We also do not find any substantial genotyping error in these variants, assuming the array is less than 99.991% accurate at sites where all samples are homozygote.

### Performance of GATK

We have run the complete “Best Practices” pipeline, including the latest version (3.6) of the “Haplotype Caller” (HC) and complete joint-calling with variant recalibration and filtering on the 97 samples^6,12^. PEMapper appears to perform as well or better than GATK in all measurable ways. GATK tends to conflate missing data with error. VCF files do not report sites that do not have high quality variant sites in at least one sample. Thus, if a site is not in the VCF file, it is not immediately clear whether the site is “missing” (insufficient evidence) or “error” (falsely believed to be high quality and reference). To try to disentangle the two in a way that displays GATK in the best possible light, we imposed the following rules. If a site was not in the VCF file, and the array called homozygous reference at the site in the sample, those sites were scored as “complete” and “agree” with the array. If a site was called variant by the array in at least one sample, but missing from the VCF file, this site was called “missing” in individuals who are not homozygous reference.

GATK calls two classes of SNPs: PASS (their highest quality calls) and Tranche99.9to100 (their second highest quality, called Tranche99.9 hereafter). Using this paradigm, GATK find theta in these samples to be .000829 (.000792 coming from PASS and .0000371 coming from Tranche99.9). GATK finds the transition-to-transversion (Ts/Tv) ratio to be 2.09 for PASS, and 1.439 for Tranche99.9, indicating that variants in Tranche99.9 are not especially trustworthy and are quite likely to be false positives.

### GATK exonic variation

GATK finds the value of theta in the exomes of these samples to be between .00041 and .00043, averaging .00423 in PASS variants. Using both PASS and Tranche99.9, theta in exomes averages .000458. The Ts/Tv ratio in exons averages 3.086 in PASS variants and 1.88 in Tranche99.9 variants. The silent-to-replacement site ratio averages 1.131 in PASS sites and 0.613 in Tranche99.9 sites, again suggesting that Tranche99.9 variants are not high quality. The individual samples averaged ~19,000 exonic variants identified by GATK PASS.

### GATK vs PEMapper/PECaller

To a great extent, PEMapper/PECaller and GATK generally make the same genotype calls at variant sites in the same samples. This is a remarkable achievement for PEMapper/PECaller, given the impressive accuracy and extensive use of training set data for GATK^32,33^. Over all 97 samples, PEMapper called 6,588,872 SNPs (single nucleotide polymorphisms with exactly two alleles) (Figure 1) with an overall transition to transversion ratio of 2.07 to 1. In category PASS there are 6,338,222 SNPs with at Ts/Tv ratio of 2.09 to 1, of these 6,241,660 (98.4%) were also called by PECaller. In Tranche99.9 there were 424,564 SNPs with a TS/TV ratio of 1.25 to 1. Of those “only” 145,373 variants were called in common with PECaller, and those SNPs had a much better Ts/Tv ratio than Tranche99.9 overall (1.72 to 1). The PASS GATK calls not made by PECaller (96,562) had a Ts/Tv ratio of 1.25 to 1. The Tranche99.9 GATK calls not made by PECaller had a Ts/Tv (266,521) ratio of 1.06 to 1. Finally PECaller SNPs not called by GATK (197,660) had a Ts/Tv ratio of 1.31 to 1. (Table 2, Figure 2) Overall, this means that PEMapper/PECaller calls slightly more variants that GATK PASS, and slightly fewer than GATK TOTAL (PASS+Tranche99.9). SNPs called by GATK, but not PEMapper/PECaller look to be of worse quality than SNPs called by PEMapper/PECaller but not GATK. The performance of Tranche99.9 SNPs in all ways suggests that they should probably not be used for analysis, as they are likely to have significant numbers of false positives.

**Table 2:** Comparison of number of variants called, and the Ts/Tv ratio for those variants, between PEMapper/PECaller and GATK. Variants not called by PEMapper/PECaller (but called by GATK) are of worse quality than those not called by GATK (but called by PEMapper/PECaller).

| **Category** | **Number of Variants Called** | **Ts/Tv Ratio** |
| --- | --- | --- |
| PEMapper/PECaller | 6,588,872 | 2.07:1 |
| GATK PASS | 6,338,222 | 2.09:1 |
| GATK 99.9 | 424,564 | 1.21:1 |
| PEMapper/PECaller and GATK 99.9 | 145,373 | 1.72:1 |
| GATK PASS but not PEMapper/PECaller | 96,562 | 1.25:1 |
| GATK 99.9 but not PEMapper/PECaller | 266,521 | 1.06:1 |
| PEMapper/PECaller but not GATK | 197,660 | 1.31:1 |

Using the Illumina 2.5M Array as the gold standard, we were able to compare the completeness and accuracy of both PEMapper/PECaller and the GATK pipeline. Across the board, PEMapper/PECaller outperformed GATK, albeit only slightly (Table 3). If the array called homozygous reference, PEMapper/PECaller was 99.95% complete and 99.94% agreed with the array, compared to GATK with 98.98% complete and 99.83% agreed. If the array called heterozygous, PEMapper/PECaller was 99.81% complete and 99.23% agreed with the array, compared to GATK with 99.31% complete and 99.78% agreed. If the array called homozygous non-reference, PEMapper/PECaller was 99.88% complete and 99.56% agreed with the array, compared to GATK with 99.68% complete and 99.15% agreed. Overall, PEMapper/PECaller was 99.85% complete and 99.76% agreed with the array, compared to GATK with 99.82% complete and 99.74% agreement with array.

**Table 3:** Comparison of calling completeness and accuracy compared to the Illumina 2.5M array gold standard for PEMapper/PECaller and GATK. PEMapper/PECaller performs slightly better than GATK.

|  | PEMapper/PECaller |  | GATK |  |
| --- | --- | --- | --- | --- |
| Variant Call Type | Completeness | Accuracy | Completeness | Accuracy |
| Homozygous Reference | 99.95% | 99.94% | 99.98% | 99.83% |
| Heterozygote | 99.81% | 99.23% | 99.31% | 99.78% |
| Homozygous Non-reference | 99.88% | 99.56% | 99.68% | 99.15% |
| Overall | 99.85% | 99.76% | 99.82% | 99.74% |

Essentially, both callers are primarily “limited” by microarray based errors. This means it may be that both callers are nearly always getting the right answer, when the array is correct, and when the array is in error, they differ, in differing ways. To a first approximation the difference between the two can be summarized, as GATK is slightly more likely than PEMapper to fail to report a site called variant by the array. The sites that GATK excludes, but PEMapper calls, are slightly more likely than average to disagree between PEMapper and the array. There is certainly no evidence that GATK is doing a substantially better job than PEMapper. We also point out that all of this is despite the fact that GATK is using knowledge about the position of high-frequency variants to help align sequences and set thresholds for calling. PEMapper/PECaller uses none of this information, and is mapping and calling variants “naively,” and yet achieves the same overall results.

In a slightly different comparison experiment, we know that with the Illumina arrays, GATK and PECaller we have three separate sets of calls. Dropping any call that is “missing” in either the array, GATK or PECaller, there are approximately 140 million genotypes called in common between the arrays and either GATK and PECaller, and over 633 million variant calls that can be compared between GATK and PECaller. For each of the three we can assume that one of the three is the “gold standard” for accuracy and ask what the error rate is at variant sites, relative to this gold standard. These results are shown in Table 4. Several conclusions can be drawn. First, all three are excellent, and in close agreement. Second, GATK looks to be a slight outlier. If GATK is set as the “gold standard,” both the array and PECaller appear to have approximately a 1% error rate at heterozygous sites, and incredibly low error rates at homozygous sites. Conversely, when comparing GATK to the array gold standard, heterozygotes appear to have an excellent error rate, but homozygous non-reference calls have an abnormally high error rate. The simplest explanation of both these observations is that GATK is slightly “over-calling” heterozygotes at the expense of homozygous calls, but only very slightly, as overall calling is truly excellent.

**Table 4:** Comparison of error rates using three possible gold standards (Illumina array, PECaller calls, GATK calls). When Illumina array calls are the gold standard, PECaller has much less error in homozygous reference and homozygous alternate calls, while having more in heterozygous calls. Overall, PECaller has slightly less error. Using all three it is possible to discern that GATK is over-calling heterozygotes at the expense of homozygous calls.

|  | Illumina Array as Gold Standard<br>(130+ million) |  | PECaller as Gold Standard |  | GATK as Gold Standard |  |
| --- | --- | --- | --- | --- | --- | --- |
|  | PECaller<br>(140 million) | GATK<br>(140 million) | Array<br>(140 million) | GATK<br>(630 million) | Array<br>(140 million) | PECaller<br>(630 million) |
| Homozygote Ref | 0.00061 | 0.00174 | 0.00224 | 0.00157 | 0.00080 | 0.00136 |
| Heterozygote | 0.00766 | 0.00217 | 0.00351 | 0.00712 | 0.01032 | 0.01132 |
| Non-ref Homozygote | 0.00439 | 0.00849 | 0.00123 | 0.00739 | 0.00107 | 0.00240 |
| All Genotypes | 0.00235 | 0.00261 | 0.00235 | 0.00300 | 0.00261 | 0.00300 |

### Insertion and deletion comparisons

Calling of insertions and deletions was not as identical as calling SNPs between the pipelines, but was still quite consistent. Overall, PECaller called 406,015 small deletions of which 84% (342,094) were called in exactly the same position by GATK. PECaller also called 212,272 insertions, of which 84% (178,478) were called by GATK. In the other direction, GATK called many more indels than PECaller. A total of 37% of deletions called by GATK, and 57% of its insertions, were not called by PECaller. This is primarily due to the fact that the Smith-Waterman mapping parameters in PEMapper were set to largely drop any read with a large (larger than ~10bp) indel. It should also be noted, that the comparison required the indel to be called in exactly the same position *i.e.* not even one base different from one another. In even slightly repetitive sequence, precise indel position is often unknowable, and it is hardly surprising that indels called by one algorithm are sometimes given slightly different positions by another. Looking at the comparison of indel genotype calls between the two pipelines, at sites called heterozygous deletions by PECaller, 98% were also called heterozygous deletions by GATK. Homozygous deletions identified by PECaller were called homozygous deletions by GATK 97% of the time. Insertions were slightly less consistent with 93% of heterozygous insertions and 94% of homozygous insertions called in common. Calling at indel sites was somewhat worse than SNPs, but still remarkably consistent, and indicative of excellent results from PEMapper/PECaller^34^.

### Exome comparison

Given that Tranche99.9 variants are of poor quality, we look at only the comparison between PEMapper/PECaller and GATK PASS variants in the exome. Overall, PEMapper/PECaller calls ~1000 more variants per exome than GATK PASS (Figure 3). The statistics for these variants are nearly identical, with PEMapper/PECaller producing a Ts/Tv ratio of 3.06 compared to 3.09 for GATK. (Figure 4) PEMapper/PECaller produced a silent/replacement ratio of 1.11 compared to 1.13 for GATK (Figure 5). Essentially, GATK appears to use its prior knowledge of variant locations to find slightly more silent sites, but may call slightly fewer potentially novel exonic replacement variants because it is limited by the existing variant lists.

### Computational time

The PEMapper and PECaller pipeline is dramatically faster than the GATK pipeline. In both cases, the first half of the pipeline was run off-site, using AWS (Amazon Web Services) resources, because the best practices requires read sorting that cannot be run in parallel on our local cluster because our cluster (like many others) uses a shared disk array environment. Total CPU time will scale similarly since all AWS instances use the same number of processors. Likewise, in both cases, the second half of the pipeline was run locally using the Emory Libraries and Information Technology’s “Tardis” resource. This computing cluster offers 12 nodes, each with 64 cores and 512GB of RAM. We report wall clock time for these tasks as well. This results in a fair comparison wherein the time to map and call 97 genomes is ~1.2 days using the PEMapper/PECaller pipeline and ~3 days using the GATK BWA/Haplotype Caller pipeline per genome analyzed (Table 5). Thus, PEMapper/PECaller is more than twice as fast even when all disk operations occured in an isolated disk environment. In a shared disk environment we could only run PEMapper. It should further be noted that PECaller jointly called the entire batch of 97 samples, something the GATK Unified Caller was incapable of doing, even on a node with 512gb of RAM. Some of the time saved using AWS is due to the fact that the GATK output is significantly larger than the output from PEMapper (approximately 150GB per sample), so the data transfer time is longer, but given that it averaged approximately over 30MB per second of transfer, this additional download time added only approximately 1.5 hours per genome. Additionally, PECaller output requires less than one tenth the data storage space as GATK. (Table 1) Including the raw sequencing data, PEMapper/PECaller requires only 40% of the storage space that GATK requires for the same sample. Finally it should be noted that since PECaller called all samples in a single batch, which allowed missing data versus homozygous reference allele calls to be distinct for all samples.

**Table 5:** Comparison of time to run PEMapper/PECaller and GATK Best Practices. PEMapper is much faster than BWA/HaplotypeCaller, while PECaller and the GATK Unified Caller and Recalibration Tool take about the same time to run. Overall, this leads to pipeline comparisons where PEMapper/PECaller is nearly twice as fast as GATK for 97 samples.

| Time to run mapping and calling |  |
| --- | --- |
| GATK (BWA and Haplotype Caller in parallel on AWS) | 70.6 +/- 9 hours per genome |
| GATK Unified Caller (batches of 10 genomes) | 0.21 hours per genome |
| GATK Recalibration Tool (batches of 10 genomes) | 0.11 hours per genome |
| **GATK Total** | **~72 hours (3 days) per genome** |
| PEMapper (in parallel on AWS) | 29 +/- 5.8 hours per genome |
| PECaller (batch of 97 genomes) | 0.34 hours per genome |
| **PEMapper/PECaller Total** | **~29.34 hours (1.2 days) per genome** |

All of this means that it is both faster and easier to run PEMapper/PECaller than the GATK pipeline for studies with more than even a handful of samples. It is also less expensive, due to the reduced usage of computational resources. Taken together, this enables more genomes to be analyzed, allowing for larger study sizes.

## Discussion

The future of genomics is WGS on 1000s+ of genomes. Analyzing that many genomes at once, both efficiently and accurately, is a tremendous computational challenge. The GATK Best Practices pipeline is the *de facto* standard for analysis of sequencing data. This is because it does an excellent job, and has proven its utility in vast numbers of exome studies. While a user may be well-advised to continue using the GATK pipeline for exome analysis, or small numbers of whole genomes^35^, we show here that PEMapper/PECaller is the decidedly better option for large-scale mapping and calling of genomes^36^. PEMapper/PECaller is significantly more efficient than GATK, requiring fewer computational resources and storage space, and thus costing less^5^.

PEMapper/PECaller manages to do this while providing nearly identical (or better) calling quality than GATK. PEMapper/PECaller also doesn’t rely on any more outside information than a reference genome, making it applicable to both human and non-human sequencing studies.

PEMapper/PECaller completely overcomes the technical challenges of GATK Best Practices. It runs well in a shared disk environment. Batch calling can occur in batches of hundreds to thousands of whole genomes easily (although computation time scales as Nlog(N) of batch size). All sites are output, together with a confidence score, so that the missing versus homozygous reference distinction is always maintained trivially. This distinction is important as it allows straightforward implementation of GWAS style QC procedures – *e.g.* sites can be filtered on call-rate and Hardy-Weinberg. The most natural way to handle these data is simply to convert them to PLINK format, QC and analyze them like any other GWAS, except that these data just happens to include all the rare and common sites from the onset.

Overall, GATK Best Practices and PEMapper/PECaller make identical calls at almost every site. When they differ from one another there is evidence that neither is very reliable. GATK Best Practices achieves its excellent results in large part by incorporating pre-existing knowledge into the pipeline. Reads are re-aligned based on preexisting knowledge of SNPs and Indels. Variants are classified, filtered, or dropped based on extensive training sets of known human variants. PEMapper/PECaller achieves essentially the same result based on no specific prior knowledge, but an intelligent genotyping model that uses nothing more than the observed data at hand. In principle, the PECaller variants could be similarly filtered/tranched/etc., but we show there is no obvious need. By not using any preexisting knowledge, PEMapper/PECaller is far easier to use in non-human systems.

PEMapper/PECaller is not only much simpler to use than GATK Best Practices, but it produces data that are of the same or very slightly higher quality. It is clear that either calling platform is more than adequate to support modern genetic studies^37^, but PEMapper/PECaller is far easier to run, uses less computational time and storage, and behaves far better in a shared disk environment. This will enable researchers to analyze large numbers of whole genomes sequences both faster and more efficiently. Using PEMapper/PECaller to map and call large-scale genome sequencing will also further precision medicine efforts^38^. Large studies utilizing whole-genome sequences are now much easier to complete computationally using PEMapper/PECaller by reducing the currently most challenging bottleneck from experiments of this type.

## ACKNOWLEDGEMENTS

We thank members of the Cutler and Zwick labs for comments on the manuscript, Cheryl T. Strauss for editing, and the Emory-Georgia Research Alliance Genome Center (EGC), supported in part by PHS Grant UL1 RR025008 from the Clinical and Translational Science Award program, the National Institutes of Health, and the National Center for Research Resources, for performing the Illumina sequencing runs. The TARDIS Emory High Performance Computing Cluster was used for this project. This work was supported by the NIH/NIMH grants U54 HD082015; U01 MH101720, which is part of the International Consortium on Brain and Behavior in 22q11.2 Deletion Syndrome (IBBC); and the Simons Foundation Autism Research Initiative [MEZ].

^#^Members of the IBBC Consortium include Bernice Morrow (Albert Einstein College of Medicine), Beverly Emanuel, Donna M McDonald-McGinn (CHOP), Steve Sharer, Anne Bassett, Eva Chow (Toronto), Joris Vermeesch, Ann Swillen (Leuven), Raquel Gur (UPenn), Carrie Bearden (UCLA), Wendy Kates (Syracuse), Vandana Shashi (Duke), Tony Simon (UC Davis), Joseph Cubells (Emory), Linda Campbell (Newcastle, Australia), Gabriela Repetto (Santiago, Chile), Jacob Vorstman (Utrecht, Netherlands), Therese Van Amelsvoort (Maastricht, Netherlands), Stephen Eliez (Geneva, Switzerland), Nicole Philip (Marseille, France), Doron Gothelf (Tel Aviv, Israel), Marianne Van Den Bree, Michael Owen (Cardiff, UK), Clodagh Murphy, Declan Murphy (London, UK), Sixto Garcia-Minaur (Madrid, Spain), Damian Neine-Suner (Mallorca, Spain), Kieran Murphy (Dublin, Ireland), Marco Armando and Stefano Vicari (Rome, Italy) as well as the named authors.

